# A low-cost pipeline for soil microbiome profiling

**DOI:** 10.1101/2020.05.07.082305

**Authors:** Anita Bollmann-Giolai, Michael Giolai, Darren Heavens, Iain Macaulay, Jacob Malone, Matthew D. Clark

**Affiliations:** John Innes Centre (JIC), Norwich Research Park, Norwich NR4 7UH, UK; Earlham Institute (EI), Norwich Research Park, Norwich, NR4 7UZ, UK; Natural History Museum (NHM), London, SW7 5BD, UK; University of East Anglia, Norwich Research Park, Norwich NR4 7TJ, UK

**Keywords:** Soil, gDNA extraction, DNA clean-up, microbiome, amplicon library construction, 16S, ITS

## Abstract

**Background:** Common bottlenecks in environmental microbiome studies are the consumable and personnel costs necessary for genomic DNA extraction and sequencing library construction. This is harder for challenging environmental samples such as soil, which is rich in PCR inhibitors. To address this, we have established a low-cost genomic DNA extraction method for inhibitor rich samples alongside an Illumina-compatible 16S and ITS rRNA gene amplicon library preparation workflow that uses common laboratory equipment. We evaluated the performance of our genomic DNA extraction method against two leading commercial soil genomic DNA kits (MoBio PowerSoil^®^ and MP Biomedicals™ FastDNA™ SPIN) and a recently published non-commercial extraction method by Zou et al. (2017). Our benchmarking experiment used four different soil types (coniferous, broad leafed, and mixed forest plus a standardised cereal crop compost mix) assessing the quality and quantity of the extracted genomic DNA by analysing sequence variants of 16S V4 and ITS rRNA amplicons.

**Results:** We found that our genomic DNA extraction method compares well to both commercially available genomic DNA extraction kits in DNA quality and quantity. The MoBio PowerSoil^®^ kit, which relies on silica column-based DNA extraction with extensive washing delivered the cleanest genomic DNA e.g. best A260:A280 and A260:A230 absorbance ratios. The MP Biomedicals™ FastDNA™ SPIN kit, which uses a large amount of binding material, yielded the most genomic DNA. Our method fits between the two commercial kits, producing both good yields and clean genomic DNA with fragment sizes of approximately 10 kb. Comparative analysis of detected amplicon sequence variants shows that our method correlates well with the two commercial kits.

**Conclusion:** Here we present a low-cost genomic DNA extraction method for inhibitor rich sample types such as soil that can be coupled to an Illumina-compatible simple two step amplicon library construction workflow for 16S V4 and ITS marker genes. Our method delivers high quality genomic DNA at a fraction of the cost of commercial kits and enables cost-effective, large scale amplicon sequencing projects. Notably our extracted gDNA molecules are long enough to be suitable for downstream techniques such as full gene sequencing or even metagenomics shotgun approaches using long reads (PacBio or Nanopore), 10x Genomics linked reads, Dovetail genomics etc.

## Background

In the last decade microbiome studies have been increasing rapidly in popularity, from 4,505 publications by December 2010 to 66,250 publications by February 2020 (PubMed reports for search term ‘microbiome’). Next generation sequencing (NGS) has made microbiome studies more accessible to a wider audience of researchers through increases in throughput and falling costs [1]. The current bottlenecks for environmental microbiome studies are the price and the hands-on time required to conduct NGS-quality genomic DNA (gDNA) extraction and NGS library preparation. Furthermore, studies sampling inhibitor rich materials such as soil [2, 3] are often restricted to the use of specialist commercial kits (costing up to £8 per extraction [4]). This cost, coupled with the many handling steps, required generally limit studies to smaller sample numbers. Although home-made gDNA extraction workflows have been described, many current methods are low in yield, throughput and are often not tested for NGS or microbiome purposes [5, 6]. Researchers face a similar bottleneck in Illumina amplicon library constructions for microbiome typing (e.g. using 16S, ITS or 18S markers). Commercial kits are limited to a small number of barcoded libraries [7], while specialist workflows (e.g. the Earth Microbiome Project benchmarked protocols) use custom sequencing primers and therefore cannot be processed using standard Illumina protocols. This limits the choice of the available sequencing provider and affects throughput and sequencing prices [8] restricting many projects to lower throughput platforms such as the Illumina MiSeq. For comparison, the Illumina MiSeq v2 500-cycle kit has an 8.5 Gb maximal output whereas the NovaSeq 6000 SP 500-cycle kit has a 400 Gb maximal output [2, 9, 10].

In scenarios with a high number of low biomass (e.g. rhizosphere samples) and inhibitor rich samples it is therefore often not feasible to perform large scale amplicon sequencing projects with a sufficient number of samples necessary for robust statistical analysis [2, 11]. To address this we developed a rapid, low cost and higher throughput <u>s</u>oil <u>D</u>NA <u>e</u>xtraction method (hereafter referred as the SDE method), for which we show equal or better gDNA extraction performance from various soil types to two leading commercial kits at a fraction of the cost per extraction (SDE: £0.2, MP Biomedicals™ FastDNA™ SPIN: £8, MoBio (now QIAGEN) PowerSoil^®^: £4). We also implemented a two-step PCR Illumina-compatible amplicon library preparation method with the capacity to multiplex 2,304 dual-barcoded sequencing libraries, at a cost of £2 per library preparation. With 438 custom designed barcodes on each end, our protocol in principle could be easily expanded to 191,844 samples per lane, sufficient for NovaSeq scale (e.g. ^~^ 10billion reads at 50,000 reads per sample). The combination of our two new protocols allows better utilisation of state of the art Illumina sequencing platforms to perform large scale amplicon-based microbiome studies at a massively reduced cost.

## Results

### DNA yield and fragment analysis of different extraction methods

We tested our SDE method by extracting gDNA from standardised (250 mg) samples of four different soil types taken from a mixed forest (MiF), a coniferous forest (CoF) and a broad-leafed forest (BrF), plus a standardised cereal crop compost mix used at the John Innes Centre (Cer). We compared our method to two leading commercial extraction kits: MP Biomedicals™ FastDNA™ SPIN and MoBio PowerSoil^®^ and a recently published low cost paperdisc method described to extract microbial DNA for PCR in less than 30 seconds [6]. We first determined which gDNA extraction method produces the highest yield and best gDNA quality. The MP Biomedicals™ FastDNA™ SPIN kit delivered the highest and the MoBio PowerSoil^®^ kit the lowest gDNA concentration (**Table 1**). The highest gDNA purity (A260/A280 and A260/A230 ratios) was obtained by the MoBio PowerSoil^®^ kit and the lowest by the MP Biomedicals™ FastDNA™ SPIN kit (**Table 1**). Our SDE method scored between the two commercial kits for both quality and quantity (**Table 1**). We further evaluated the methods for the extracted gDNA fragment length. For the MoBio PowerSoil^®^ kit method we found fragment lengths between 13.9 kb and 24.4 kb. The SDE method produced fragments between 11.3 kb and 11.7 kb and the MP Biomedicals™ FastDNA™ SPIN mostly extracted fragments below 10 kb (**Additional File 1**). For the paperdisc method we could only measure a gDNA concentration as there was insufficient DNA for fragment analysis.

**Table 1.** Genomic DNA quality control: Qubit BR DNA kit was used to measure the total gDNA concentration and the nanodrop was used to compare the A260/280 and A260/230 ratios. The MoBio PowerSoil^®^ kit delivers high quality but low yields whereas the MP Biomedicals™ FastDNA™ SPIN kit delivers high yields at low quality. The SDE method falls between the two commercial kits for both quality and quantity. CoF stands for coniferous forest, MiF for mixed forest, BrF for broad leafed forest and Cer for John Innes cereal compost mix.

| Soil type | MoBio |  |  | MP |  |  | SDE |  |  |
| --- | --- | --- | --- | --- | --- | --- | --- | --- | --- |
|  | [ng] | A260/280 | A260/230 | [ng] | A260/280 | A260/230 | [ng] | A260/280 | A260/230 |
| Cer | 0 ± 0 | -8.77 ± 18.65 | 0.82 ± 0.3 | 1914.67 ± 57.87 | 1.75 ± 0.07 | 0.06 ± 0.01 | 468 ± 26.46 | 1.31 ± 0.05 | 0.74 ± 0.02 |
| MiF | 1203.33 ± 540.12 | 2.04 ± 0.19 | 1.77 ± 0.46 | 20333.33 ± 3395.13 | 1.87 ± 0.02 | 0.26 ± 0.04 | 2913.33 ± 451.81 | 1.45 ± 0.07 | 0.86 ± 0.1 |
| BrF | 1036 ± 699.6 | 2.12 ± 0.09 | 1.68 ± 0.3 | 8400 ± 1805.33 | 1.82 ± 0.06 | 0.16 ± 0.04 | 1810 ± 671.79 | 1.33 ± 0.02 | 0.87 ± 0.02 |
| CoF | 73.33 ± 127.02 | 2.02 ± 0.59 | 0.90 ± 0.38 | 6920 ± 682.35 | 1.26 ± 0.4 | 0.35 ± 0.28 | 942.67 ± 241.17 | 1.29 ± 0.01 | 0.60 ± 0.01 |

### Extraction method effects on bacterial and fungal amplicon library construction

We constructed 16S V4 and ITS rRNA Illumina sequencing libraries from all extractions (3 biological replicates per soil type) using 3 ng of gDNA input per library construction reaction and three technical replicates (similar to [12]). All libraries were quality controlled using LabChip GX Touch high sensitivity capillary electrophoresis. The gDNA extracted with MP Biomedicals™ FastDNA™ SPIN, MoBio PowerSoil^®^ and our SDE method performed well in library construction, producing libraries with similar profiles (**Figure 2**). The paperdisc method did not produce a library with a detectable electropherogram trace. We pooled all libraries at equal mass (except the paperdisc method where we used the maximal amount as these libraries were not detectable) and submitted each library for 250bp paired-end sequencing. Sequencing of the paperdisc extraction method did not produce any reads suggesting that the extracted gDNA concentration was too low for successful library construction.

**Figure 1.**
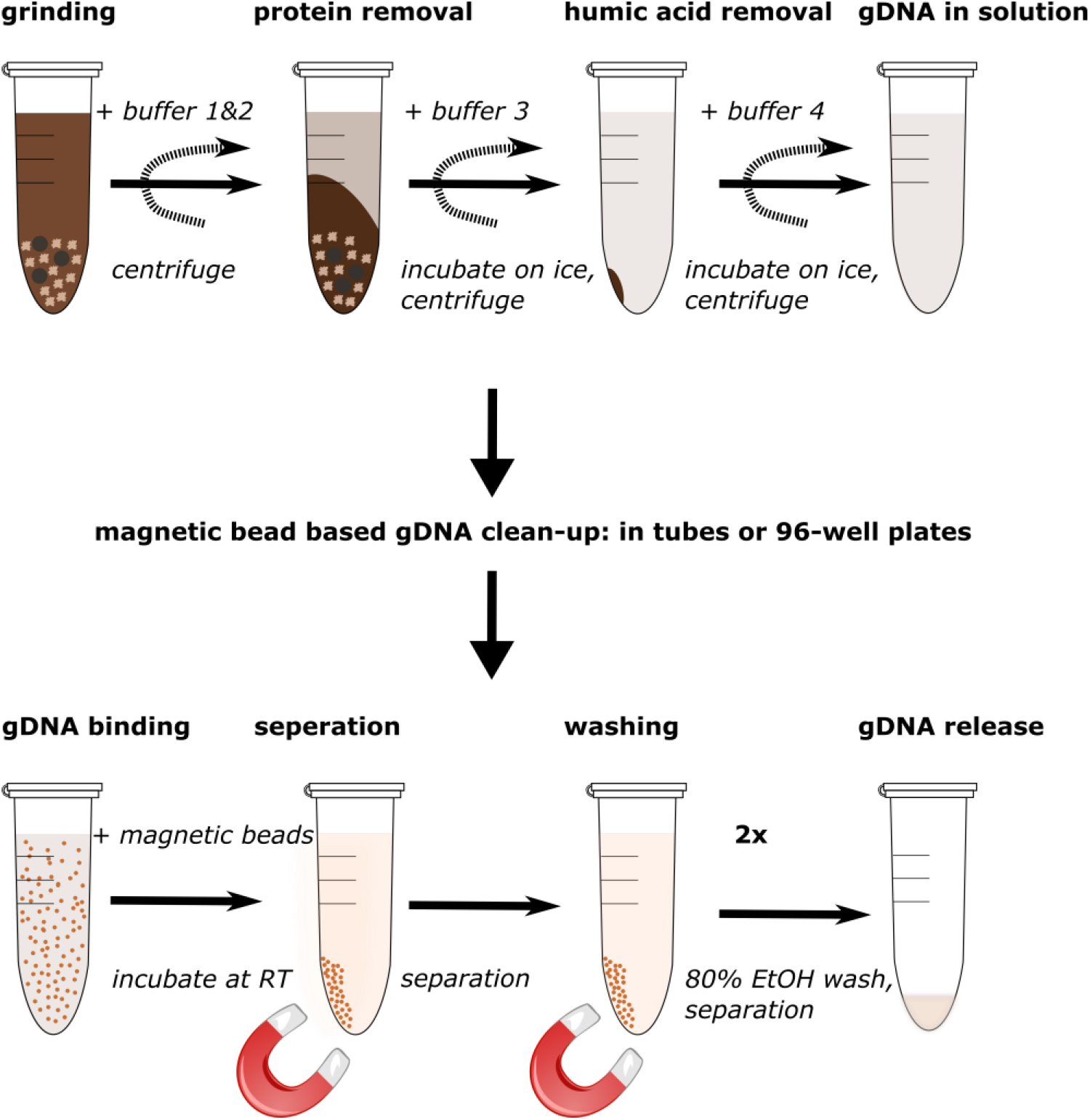
Overview of the SDE soil gDNA extraction protocol: The genomic DNA extraction protocol is divided in two parts, the gDNA extraction and the gDNA clean-up. The soil material is grinded with buffer one and two, following centrifugation the proteins are removed with buffer three and incubation on ice. After centrifugation, humic acid is removed by using buffer four and incubation on ice. The extracted genomic DNA is then ready for clean-up after centrifugation. The gDNA clean-up is performed using magnetic bead-based isopropanol precipitation of the gDNA, plus two bead washing steps with 80% Ethanol. The gDNA may then be eluted into a buffer of choice – in our case TE.

**Figure 2.**
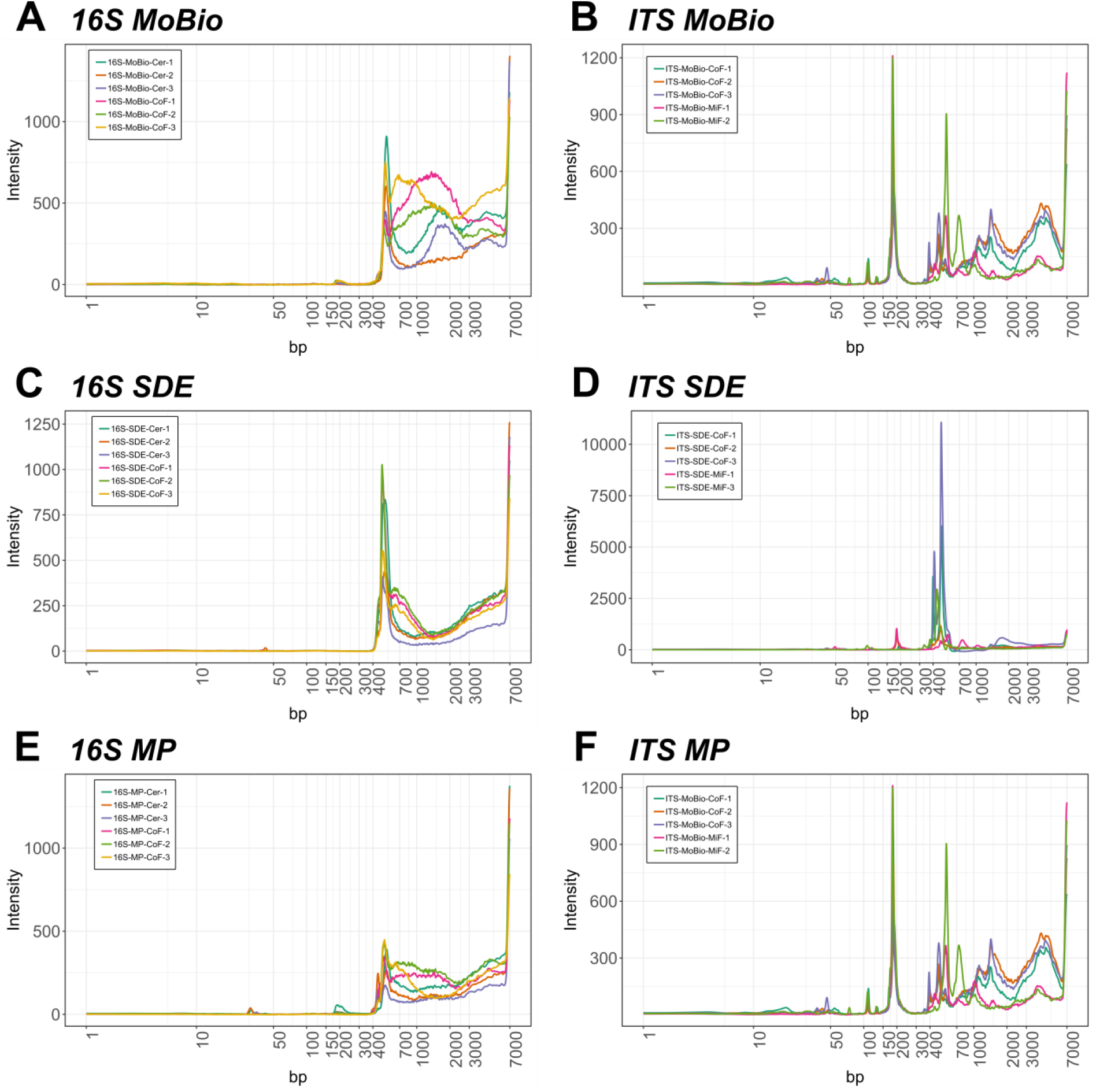
Representative results comparing MoBio PowerSoil^®^, MP Biomedicals™ FastDNA™ SPIN and SDE methods: (**A**) Representative 16S libraries of gDNA extracted with MoBio PowerSoil^®^. (**B**) Representative ITS libraries of gDNA extracted with MoBio PowerSoil^®^. (**C**) Representative 16S libraries of gDNA extracted with SDE. (**D**) Representative ITS libraries of gDNA extracted with SDE. (**E**) Representative 16S libraries of extracted gDNA with MP Biomedicals™ FastDNA™ SPIN. (**F**) Representative ITS libraries of extracted gDNA with MP Biomedicals™ FastDNA™ SPIN. Target size of representative 16S libraries is between 350bp and 500bp and for ITS libraries between 300bp and 700bp.

### Comparison of extraction methods based on bacterial and fungal microbial composition

It has been previously reported that different microbial gDNA extraction methods can introduce a genera bias [13]. To test this, we compared the biological replicates of each library preparation method (apart from the paperdisc method) by correlation analysis of the detected bacterial and fungal Amplicon Sequence Variants (ASVs) (**Additional File 2**) The three tested library construction methods compared well across all soil types. We also compared the genus abundances of our SDE method with the two commercial kits by analysing the bacterial and fungal genus abundances of each library. The genus abundance plots for each soil type were not statistically significantly different between the extraction methods used for either fungal (adonis test, p-value: 0.787) or bacterial communities (adonis test, p-value:0.603). Instead our data showed statistically significant variation between soil types (bacteria ANOSIM test, p-value: 9.99e-4, fungi ANOSIM test, p-value: 9.99e-4) but not between gDNA extraction methods (bacteria: **Figure 3A**, **Figure 3B** and fungi: **Figure 3C, Figure 3D**). We further tested the samples using beta-diversity as a measure (Bray-Curtis) for between-sample similarity. This analysis agreed with the result of the genus abundance plots i.e. by clustering the soil types separately but not clustering the data for the three extraction methods (**Figure 3B**, **Figure 3D**). We confirmed this result with permutation multivariate analysis of variance analyses (PERMANOVA, package ‘vegan’ version 2.5.2, adonis function).

**Figure 3.**
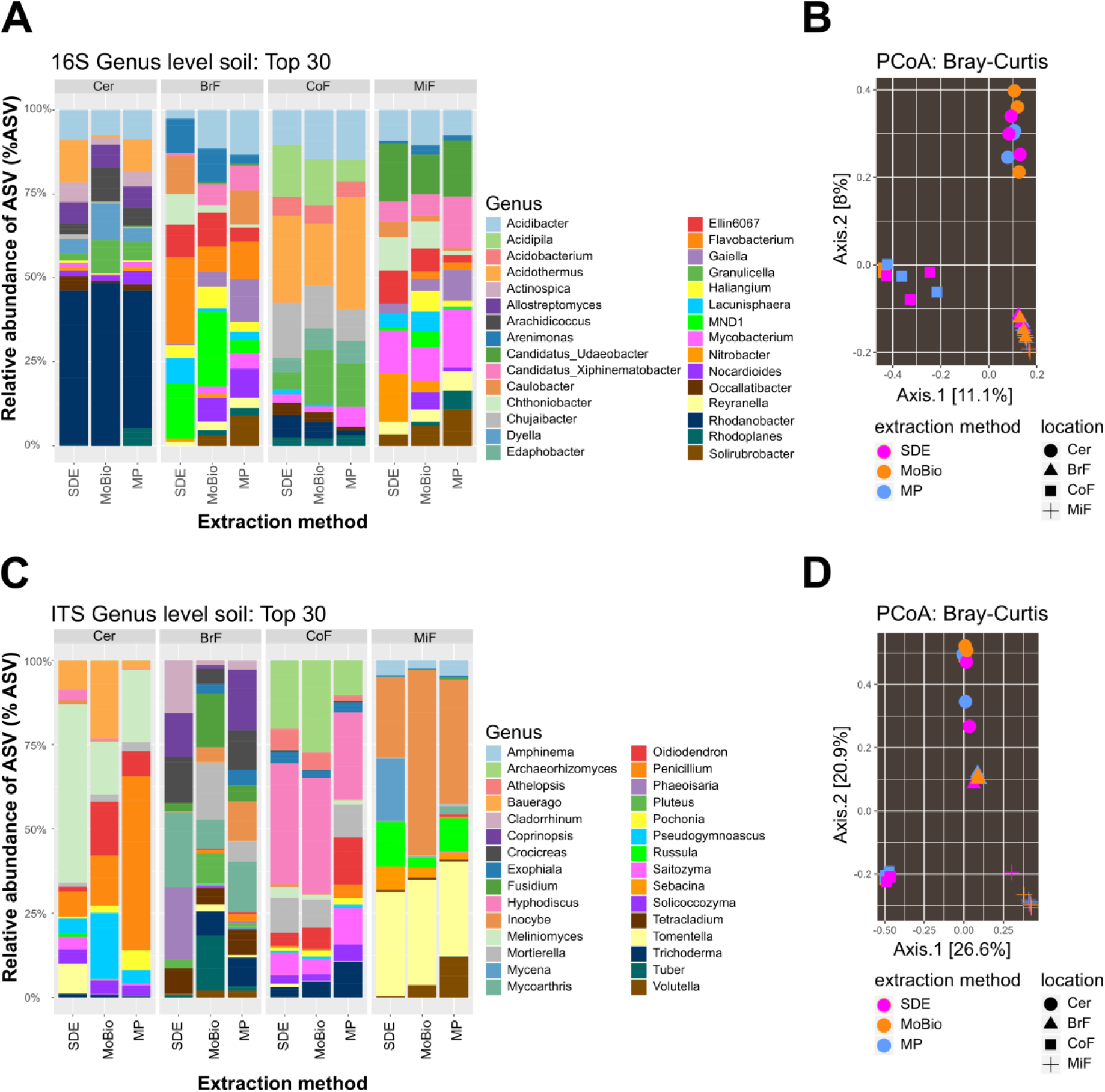
PCoA comparison of ASV abundances and relative abundance bar chart for bacterial and fungal communities of three different gDNA extraction methods and four soil types: (**A**) Top 30 bacterial community composition representing 0.72% of the overall bacterial community. (**B**) PCoA showing beta-diversity by Bray-Curtis distance for bacterial community composition. (**C**) Top 30 fungal community composition representing 1.87% of the overall fungal community. (**D**) PCoA showing beta-diversity by Bray-Curtis distance for fungal community composition. **A** and **B** show that the clustering for the entire bacterial communities is soil-type dependant and not driven by different gDNA extraction methods. **C** and **D** show the same is true for fungal community structure. Cer represents a standard cereal compost used at the JIC, Norwich, UK; BrF represents a broad-leafed forest soil; CoF represents a coniferous forest soil and MiF represents a mixed forest soil. Bead represents the SDE gDNA extraction method, MoBio represents MoBio PowerSoil^®^ and MP represents MP Biomedicals™ FastDNA™ SPIN. Statistical analysis shows no significant difference between DNA extraction methods used for both bacterial (ANOSIM test, p-value:0.603) and fungal (ANOSIM test, p-value:0.787) communities, but shows significant difference between locations for bacterial (adonis test, p-value: 9.99e-4) and fungal (adonis test, p-value: 9.99e-4) communities.

To compare the extraction methods in more detail we correlated the detected ASVs abundances of each kit. In the SDE to MoBio PowerSoil^®^ kit comparison we found the following correlation coefficients for bacterial ASVs: Cer 0.75, BrF 0.74, CoF 0.94, MiF 0.85 (**Figure 4A**) and for fungal ASVs: Cer 0.88, BrF 0.49, CoF 0.85, MiF 0.86 (**Figure 4B**). The correlation analysis of the SDE to MP Biomedicals™ FastDNA™ SPIN kit delivered similar results (Cer 0.84, BrF 0.6, CoF 0.9, MiF 0.73 for bacteria, **Additional File 3A** and Cer 0.8, BrF 0.49, CoF 0.82, MiF 0.82 for fungi, **Additional File 3C**). These results confirm that our SDE method fits between two commonly used commercial soil gDNA extraction kits across a broad range of measurable parameters.

**Figure 4.**
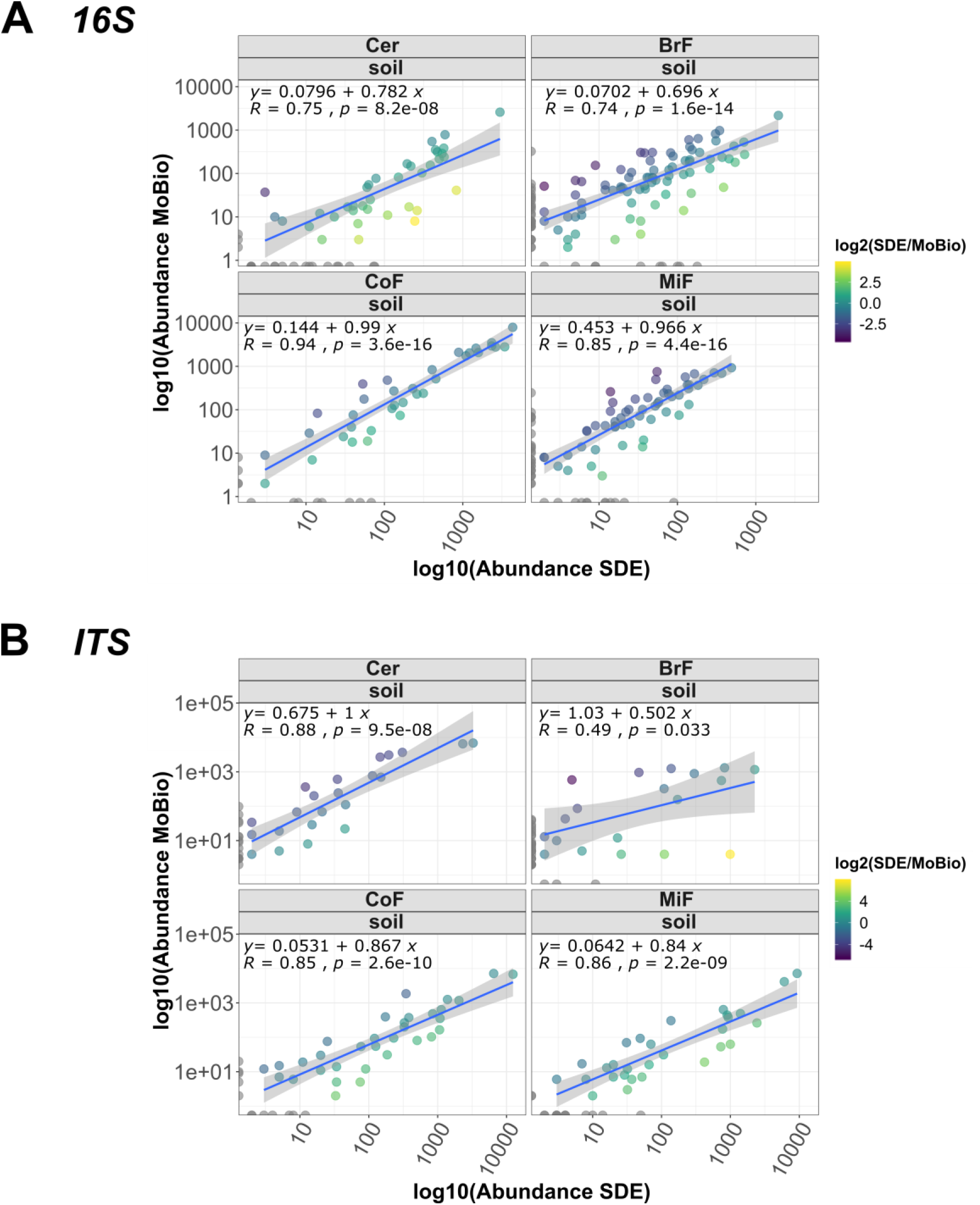
Correlation analysis of bacterial and fungal ASVs from SDE versus MoBio PowerSoil^®^ gDNA extraction: (**A**) shows bacterial ASV correlation between MoBio PowerSoil^®^ and SDE for four different soil types. (**B**) shows fungal ASV correlation between MoBio PowerSoil^®^ and SDE for four different soil types. Cer: Standard cereal crop soil mix used at JIC, Norwich, UK. BrF: soil from a broad-leafed forest. CoF: soil from a coniferous forest. MiF: Soil from a mixed forest. Both extraction methods led to bacterial and fungal ASV that show positive correlation, indicating that there is no extraction method-induced bias regarding sequenced ASV. Abundances on x and y axes were log10 scaled, each dot represents a genus, dots are coloured as log2(Abundance(SDE)/Abundance(Commercial Kit)). Grey dots on each axis are Genera uniquely detected for the extraction method.

## Discussion

Custom (non-kit) gDNA extraction methods for inhibitor rich samples have been described previously [14–19]. These methods emphasised gDNA quantity [18, 19], quality [14, 18, 19] or cost-efficiency [6, 17, 18, 20]. However, an often overlooked but practically important consideration is the hands-on time required per extraction without a quantity or quality penalty. KatharoSeq [21] for example is a pipeline for low biomass samples that delivers good gDNA quality with less hands on time, however, it still uses parts of a commercial kit, which increases the price per sample. On the other hand Zou et al. described a fast and very affordable gDNA extraction method but at the cost of poor gDNA quantity [6]. Here we present a high-throughput gDNA extraction method that is suitable for low input and inhibitor rich sample types such as soils (**Figure 1**). We compared our protocol to two leading commercial (MoBio PowerSoil^®^ and MP Biomedicals™ FastDNA™ SPIN kits) and a non-commercial (paperdisc based) extraction method [6] and observed that our SDE method delivered good quality and quantities of NGS-compatible gDNA at a fraction of the cost of commercially available solutions. SPRI-bead-based methods allow fast, scalable and inexpensive extraction of nucleic acids [22]. We designed our extraction method to be compatible with 96-well format plates. Because it uses simple pipetting steps, SPRI-bead based purification and washing steps, our method should be easily adaptable from multi-channel pipette to common liquid handling robotic systems. The extracted gDNA from four distinctively different soil types using SDE is similar in quality and quantity to the two commercial kits (**Table 1**), with the extracted gDNA from the commercial kits and the SDE method led to similar amplicon library profiles (**Figure 2**). In contrast, the paperdisc method did not generate useable sequencing libraries. We further investigated the gDNA preparation methods for extraction biases when analysing fungal and bacterial communities. Here, the results of the two commercial kits and the SDE method overlap, showing no statistical differences in microbiome composition (**Figure 3**). Clustering of the sequencing data using PCoA separated soil types but not gDNA extraction methods (**Figure 3B, Figure 3D**). A correlation analysis between SDE and MoBio, SDE and MP, MoBio and MP, for detected bacterial and fungal communities (**Figure 4**, **Additional File 2**, **Additional File 3**) showed strong correlation between the commercial kits and the SDE method. This altogether indicates that our SDE method does not induce an experimental bias in extracting bacterial and fungal community data.

To conclude, we present a low-cost gDNA extraction method (£0.2/sample) that effectively extracts inhibitor-rich samples and delivers good quality and quantities of gDNA suitable for microbiome studies. We show that the SDE method does not introduce a library preparation and therefore a sequencing bias. We also present a low-cost custom and fully Illumina compatible bacterial and fungal amplicon library construction protocol that can multiplex up to 2,304 samples to one pool (£2/library). Our method therefore enables researchers to sequence their projects on any available Illumina platform without the need to purchase full lanes/flow cells. Overall, the two presented methods will enable microbiome projects to be performed at any desired scale at an affordable price for a broad audience of microbiome enthusiasts.

## Methods

### Soil material collection

We selected three different woodlands for soil collection: a coniferous forest (52.661750, 1.095444, UK), a mixed forest (46.394474, 11.235371, Italy) and a broad-leafed forest (46.454682, 11.301284, Italy). We sampled the soil material from the topsoil into sterile 50 ml conical tubes (Supplier Starlab (UK) Ltd, E1450-0800) using nitrile gloves and a sterilised shovel. The sampled material was stored in a mobile refrigerator during transportation to the lab where the soil was stored at 4 °C until use for gDNA extraction. The cereal compost mix was collected in the same manner but obtained from the John Innes Horticulture facilities (Norwich, U.K.).

### MoBio PowerSoil^®^ DNA Isolation Kit

The kit was applied following to the manufacturer’s instructions with one alteration. The soil material was ground using the Geno/Grinder (SPEX SamplePrep 2010) for one minute at 1,750 rpm using the supplied grinding stones. The active hands-on time without incubation and centrifugation times is 17 min per extraction.

### MP Biomedicals™ FastDNA™ SPIN Kit for Soil

The kit was applied following to the manufacturer’s instructions with no alterations, using the recommended Fastprep machine (Fastprep24, MP BIO) for the grinding step. The active hands on time without incubation and centrifugation times is 10 min per extraction.

### Paperdisc method

The paperdisc extraction used is based on the protocol by Zou et al. 2017 [6]. We followed the protocol as described previously [6] with one alteration: instead of using one paperdisc (Whatman qualitative filter paper, WHA1001070, SIGMA-ALDRICH CO LTD) per extraction, in an attempt to increase yields, we added four discs to the washing buffer. We transferred a disc to 50 μl TE buffer (10 mM Tris-HCl pH 8.0, 1 mM EDTA) for four hours and measured the gDNA concentration using a Qubit 2.0 Fluorometer (Thermo Fisher Scientific, Waltham, USA) with Qubit dsDNA BR DNA assay kit reagents (Q32853, Thermo Fisher Scientific, Waltham, USA).

### SDE method

250 mg soil was transferred to a 2 ml tube containing 300 μl sterile 1 mm diameter garnet particles (STRATECH SCIENTIFIC LTD, 11079110gar-BSP, Biospec products). Before grinding we added 750 μl Buffer 1 (181 mM Trisodium phosphate, 121 mM guanidinium thiocyanate, 0.22 μM sterile filtered with Sartorius UK Ltd, 16532K) and 60 μl Buffer 2 (150 mM NaCl, 4% SDS, 0.5M Tris pH7, 0.22 μM sterile filtered) to each tube. To lyse the bacterial and fungal cells the tubes were transferred to the Genogrinder (Spex SamplePrep, 2010) and run for 1 minute at 1,750 rpm. We centrifuged the tubes at 17,000 rcf for 2 minutes to pellet debris, and 450 μl supernatant were transferred to a new 2 ml tube, mixed with 250 μl Buffer 3 (133 mM Ammonium acetate, 0.22 μM sterile filtered) and incubated for 10 minutes on ice to precipitate proteins. We centrifuged the tubes at 17,000 rcf for 3 minutes and transferred 500 μl clear supernatant to a new 2 ml tube. The supernatant was mixed with 200 μl Buffer 4 (60 mM aluminium sulphate, 0.22 μM sterile filtered) and the reaction incubated for 10 minutes on ice to enable additional protein precipitation and humic acid removal. We centrifuged the reaction at 17,000 rcf for 10 minutes and transferred either 600 μl supernatant to a 1.5 ml tube or 140 μl supernatant to a 96 well plate. Supernatants were stored at −20°C until further use. Buffer composition for Buffer 1, 2, 3 and 4 adapted from Brolaski et al. [23].

### SDE single tube gDNA clean-up

To perform gDNA clean-ups in single tubes we prepared 10 μl magnetic beads (Sera-Mag Carboxylate-Modified Magnetic Particles (Hydrophophylic), 24152105050250, GE Healthcare Life Sciences) per extraction. The beads were transferred to a 2 ml tube and washed twice with a large volume of in 1 % Tween-20. After washing we eluted 10 μl beads in 20 μl 1 % Tween-20 and added 20 μl of the Tween-20/bead mixture to each extraction (a final Tween-20 concentration of ^~^ 0.02%). DNA was precipitated onto the beads for 5 – 10 minutes by adding 0.7 × volume of Isopropanol (420 μl). We washed the magnetic beads twice with 500 μl 80% ethanol on a magnet rack, air dried the beads and eluted the gDNA in 50 μl 1 × TE buffer (10 mM Tris-HCl pH 8.0, 1 mM EDTA) for 5 – 10 minutes. The eluted gDNA was transferred to a 1.5 ml tube and stored at −20°C until further use.

The active hands on time without centrifugation and incubation times is 8 min per extraction.

### SDE 96 well plate gDNA clean-up

To perform the gDNA clean-ups in a 96-well format we transferred 140 μl supernatant to a 96 well plate (96 Well Non-Skirted PCR Plate, CLEAR, 4TI-0750_50, 4titude Ltd, UK). For each extraction we washed 5 μl magnetic beads and eluted the beads in 5 μl 1 % Tween-20 (for 96 samples we therefore prepared 480 μl magnetic beads). We added 5 μl of the 1 % Tween-20 / bead mixture to each reaction and precipitated the DNA on the beads using a 0.7 × volume Isopropanol (98 μl). We mixed the reactions by vortexing for 5 seconds and incubated the plates for 5 minutes to precipitate the gDNA. We then washed the beads twice on a magnet rack with 100 μl 80% ethanol, removed the remaining ethanol well and eluted the gDNA in 50 μl 1 × TE buffer for 10 minutes. The cleaned gDNA was transferred to a fresh 96 well plate (96 Well Non-Skirted PCR Plate, CLEAR, 4TI-0750_50, 4titude Ltd, UK) and stored at −20 °C until further use.

### Genomic soil DNA extraction and quality control

We performed the gDNA extraction using three replicates per soil sample and extraction method, using 250 mg soil from the same sample for each extraction. We tested four different extraction methods: (a) the PowerSoil^®^ DNA Isolation Kit (12888-50, CAMBIO, UK, now 12888-100, DNeasy PowerSoil, QIAGEN LTD), (b) the MP Biomedicals™ FastDNA™ SPIN Kit for Soil (11492400, Fisher scientific, UK), (c) our SDE method and (d) the recently published paperdisc method [6].

We determined the gDNA concentrations (**Table 1**) using the Qubit 2.0 Fluorometer (Thermo Fisher Scientific, Waltham, USA) dsDNA BR DNA assay kit (Q32853, Thermo Fisher Scientific, Waltham, USA) and assessed the purity of the extracted gDNA by measuring the 260:280 and 260:230 absorbance ratios using the NanoDrop ND-1000 Spectrophotometer (**Table 1**). To assess the fragment size of the extracted gDNA we used the 2200 TapeStation (Agilent Technologies, Stockport, UK) genomic DNA screen tape (5067-5365, Agilent Technologies, Stockport, UK) and genomic DNA reagents (5067-5366, Agilent Technologies, Stockport, UK) (**Figure 2A**).

### Amplicon library construction, quality control and pooling

We targeted the bacterial variable (V4) region of the 16S rRNA gene and the ITS1 region of the Internal Transcribed Spacer (between 18S and 5.8S rRNA subunit) for amplicon library construction. Amplicon library construction was performed using a two-step PCR protocol: In a first PCR we targeted the 16S V4 and ITS regions using gene specific primers with a 5’ primer tail that allows the addition of the barcoded sequencing adapters for custom dual indexing in a second PCR [24, 25] (**Additional File 4**). For the bacterial 16S V4 PCRs we included PNAs (peptide nucleic acid PCR clamps) targeting plant mitochondria (mPNA) and chloroplast regions (cPNA) [26] in our first PCR to minimise plant gene amplification.

For the first PCR we used the following primers adapted from Walters et al [8]: 16S 515 forward 5’-[TCGTCGGCAGCGTC][AGATGTGTATAAGAGACAG][GT][GTGYCAGCMGCCGCGGTAA]-3’ (5’-[P5][Tn5 adapter][linker][16SV4]-3’), 16S 806 reverse 5’-[GTCTCGTGGGCTCGG][AGATGTGTATAAGAGACAG][CC][GGACTACNVGGGTWTCTAAT]-3’ (5’-[P7][Tn5 adapter][linker][16SV4]-3’), ITS1 forward 5’-[TCGTCGGCAGCGTC][AGATGTGTATAAGAGACAG][GG][CTTGGTCATTTAGAGGAAGTAA]-3’ (5’-[P5][Tn5 adapter][linker][ITS]-3’) and ITS2 reverse 5’-[GTCTCGTGGGCTCGG][AGATGTGTATAAGAGACAG][CG][GCTGCGTTCTTCATCGATGC]-3’ (5’-[P7][Tn5 adapter][linker][ITS]-3’). All primers were ordered from Integrated DNA Technologies (IDT, Leuven, Belgium). The 5’ tails of the gene 16S V4 and ITS specific primers contain the Illumina Nextera Tn5 transposase adapter and linker sequences. This allows further amplification of the amplicons with barcoded Illumina adapters and sequencing using Illumina chemistry with the Illumina supplied sequencing or indexing primers (all oligonucleotide sequences are in **Additional File 4**).

We performed the first PCR step using 3 ng gDNA input, 1 Unit Kapa HiFi polymerase (KK2102, Roche, UK), 1x Kapa HiFi Fidelity buffer (KK2102, Roche, UK), 0.25 μM reverse and 0.25 μM forward primer (IDT, UK), 1x KAPA Enhancer 1 (KK5024, SIGMA-ALDRICH CO LTD, UK), 0.3 μM dNTPs (KK2102, Roche, UK), 1.25 μM cPNA (PNA BIO INC, USA), 1 μM mPNA (PNA BIO INC, USA) and DNase/RNase free distilled water (10977-049, Thermo Fisher Scientific, UK) in 10 μl total reaction volume. We performed each PCR with three technical replicates using the following cycle conditions: initial denaturation at 95 °C for 3 minutes, followed by 20 cycles of denaturation at 98 °C for 20 seconds, PNA clamping at 75 °C for 10 seconds, primer annealing at 55 °C for 30 seconds, elongation at 72 °C for 30 seconds, with a final elongation step at 72 °C for 3 minutes (Alpha Cycler 4, PCRmax, Labtech International Ldt., UK). The PCR mix for the fungal libraries was the same but with ITS instead of 16S V4 primers and without the PNA oligo blockers. ITS amplification reactions were run with the following cycle conditions: initial denaturation at 95 °C for 3 minutes, followed by 20 cycles of denaturation at 98 °C for 20 seconds, annealing at 55 °C for 30 seconds, elongation at 72 °C for 30 seconds, with a final elongation at 72 °C for 3 minutes (Alpha Cycler 4, PCRmax, Labtech International Ldt., UK).

After this first gene targeting PCR step, we pooled the three technical replicates reactions for the same sample and conducted a 0.7x magnetic bead clean-up (HighPrep™ PCR Clean-up System, AC-60050, MAGBIO). The clean PCR products were eluted in 10 μl 1 × TE buffer. The second PCR was conduct on the pooled replicate sample with the same program for bacterial and fungal libraries i.e. using 1x Kapa HiFi Fidelity buffer (KK2102, Roche, UK), 1 Unit Kapa HiFi polymerase (KK2102, Roche, UK), 0.2 μM P5 indexing primer (IDT, Leuven, Belgium), 0.2 μM P7 indexing primer (IDT, Leuven, Belgium), 0.3 μM dNTPs (KK2102, Roche, UK), 7.6 μl of clean gene targeting PCR product and DNase/RNase free distilled water (10977-049, Thermo Fisher Scientific, UK) in a total reaction volume of 30 μl. The barcoding cycle conditions (Alpha Cycler 4, PCRmax, Labtech International Ldt., UK) were: initial denaturation at 95 °C for 3 minutes, followed by 15 cycles of denaturation at 98 °C for 20 sec, annealing at 62 °C for 30 sec, elongation at 72 °C for 30 sec, with a final elongation at 72 °C for 3 minutes. After the barcoding PCR step the reactions were cleaned using a 0.7x magnetic bead clean-up (HighPrep™ PCR Clean-up System, AC-60050, MAGBIO) and the final libraries eluted in 20 μl EB buffer (10 mM Tris-HCl pH 8.5).

We quantified the cleaned amplicon libraries using the Qubit 2.0 Fluorometer (Thermo Fisher Scientific, Waltham, USA) with dsDNA HS Assay Kit reagents (Q32854, Thermo Fisher Scientific, Waltham, USA) and controlled the size of the amplicons on the GX Touch using the 3K kit (X-Mark DNA LabChip, CLS144006, HT DNA NGS 3K Reagent Kit, Perkin Elmer LAS (UK) LTD). The amplicons were pooled equimolarly to 1.5 nM for bacterial and 2.0 nM for fungal libraries according to the molarity obtained by the LabChip GX Touch smear analysis of the region between 380 and 650 bp. We performed a final 0.7 × magnetic bead clean-up (HighPrep™ PCR Clean-up System, AC-60050, MAGBIO) of the libraries and eluted the final library pools in 50 μl EB buffer.

### Sequencing

The 16S and ITS library pools were sequenced using the MiSeq Nano reagent version 2, 500 cycle kit (Illumina, UK) at an 8 pM loading concentration with a 10 % PhiX spike-in. 16S and ITS pools were sequenced separately – each pool using a MiSeq Nano reagent kit.

### Amplicon data analysis

We demultiplexed bcl files using bcl2fastq version 2-2.20.0.422 with the settings --barcode-mismatches 1 --fastq-compression-level 9 into individual fastq.gz files. We trimmed the paired-end reads for primers, sequencing adapters and linker sequences using cutadapt-1.9.1 [27] with the settings -n 4 --minimum-length=50. The data was quality controlled using R-3.5.0 and DADA2 version 1.8.0 according to the workflow described in [28] version 2. The truncation length for forward reads was set to 180 bp and the truncation length for the reverse reads to 200 bp. For 16S libraries we used the following parameters: maxN=0, maxEE=2 and truncQ=11. For ITS libraries we specified the following parameters: maxN=0, maxEE=c(2, 2) and truncQ=11 and a minimum length of 50 bp. Forward and reverse reads were merged with default settings. We used the Silva (silva_nr_v132) database to classify bacterial reads [29] and UNITE (sh_general_release_dynamic_s_01.12.2017) for the fungal dataset [30]. Reads that did not match to the bacterial or fungal database and ASVs with a mean lower as 10^−5^ were removed from the datasets. The filtered data with 4,113 bacterial ASVs and 1,602 fungal ASVs (package ‘phyloseq’, version 1.24.0) was used to calculate the β-diversity (Bray-Curtis, R-3.5.0 ‘vegan’ package, version 2.5.2) and to perform statistical analysis (package ‘vegan’, ANOSIM and PERMANOVA: adonis function) [31]. All numbers of processed reads through the analysis pipeline are in **Additional File 5**.

We performed the correlation analysis in R-3.5.0 using the filtered phyloseq object on genus level and plotted it with log10 scaling [32]. The corrplot was generated in R, using the filtered phyloseq object on order level with the corrplot package version 0.84. All figures were generated in R-3.5.0 using the R package ggplot2-3.1.1 [33].

## Additional files

Additional File 1 (Word file): Supplemental figures of genomic tapestation traces of representative MoBio PowerSoil^®^, MP Biomedicals™ FastDNA™ SPIN and SDE gDNA extractions.

Additional File 2 (PDF file): Corrplot of technical replicates of MP Biomedicals™ FastDNA™ SPIN, MoBio PowerSoil^®^ and SDE method. Order level.

Additional File 3 (PDF file): Correlation analysis SDE method versus MP Biomedicals™ FastDNA™ SPIN & MP Biomedicals™ FastDNA™ SPIN vs MoBio PowerSoil^®^(16S/ITS)

Additional File 4 (Excel file): List of primers used for library construction and barcodes.

Additional File 5 (Excel file): read numbers through QC and analysis pipeline steps.

## Availability of data and materials

### ENA

Sequencing reads for the all experiments are available under the ENA study Accession Number PRJEB37921.

## List of abbreviations

ASVs: amplicon sequence variants
BrF: broad leafed forest
Cer: John Innes cereal compost mix
CoF: coniferous forest
gDNA: genomic DNA
ITS: internal transcribed spacer
MiF: mixed forest
NGS: Next generation sequencing
PNA: peptide nucleic acid
SDE: soil DNA extraction

## Declarations

Not applicable.

## Ethics approval and consent to participate

Not applicable.

## Consent for publication

All authors give consent for publication.

## Competing Interests

The authors have no competing interests.

## Funding

Financial support was provided by a John Innes Foundation Rotation PhD Studentship to A.B.G and Natural History Museum support to M.D.C. Work in the Malone lab is funded by Biotechnology and Biological Sciences Research Council (BBSRC) Institute Strategic Programme grant BBS/E/J/000PR9797 (Plant Health) to the John Innes Centre.

## Authors’ contributions

A.B.G established methods, analysed the data and wrote the manuscript. M.G. contributed to method development and data analysis. D.H. helped with gDNA quality controls. M.D.C designed the barcode sequences. I.M., J.M. and M.D.C helped write the manuscript. A.B.G and M.D.C designed the study.

## Acknowledgements

We thank Sarah Walkington at the Natural History Museum, London for assistance with sequencing.

## Notes

### Competing Interest Statement

The authors have declared no competing interest.

