## Additional_File_1 for "A low-cost pipeline for soil microbiome profiling"

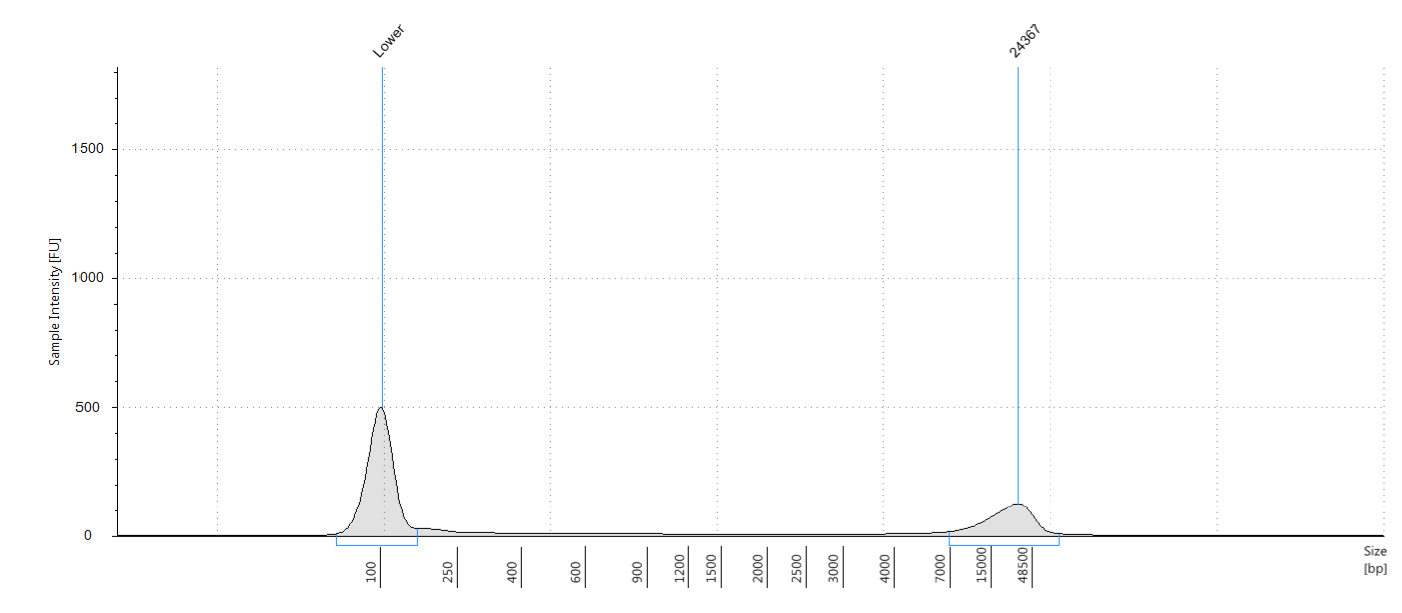

**Figure 1 Tapestation gDNA trace MoBio PowerSoil® for representative Cer (John Innes cereal compost mix) sample.**

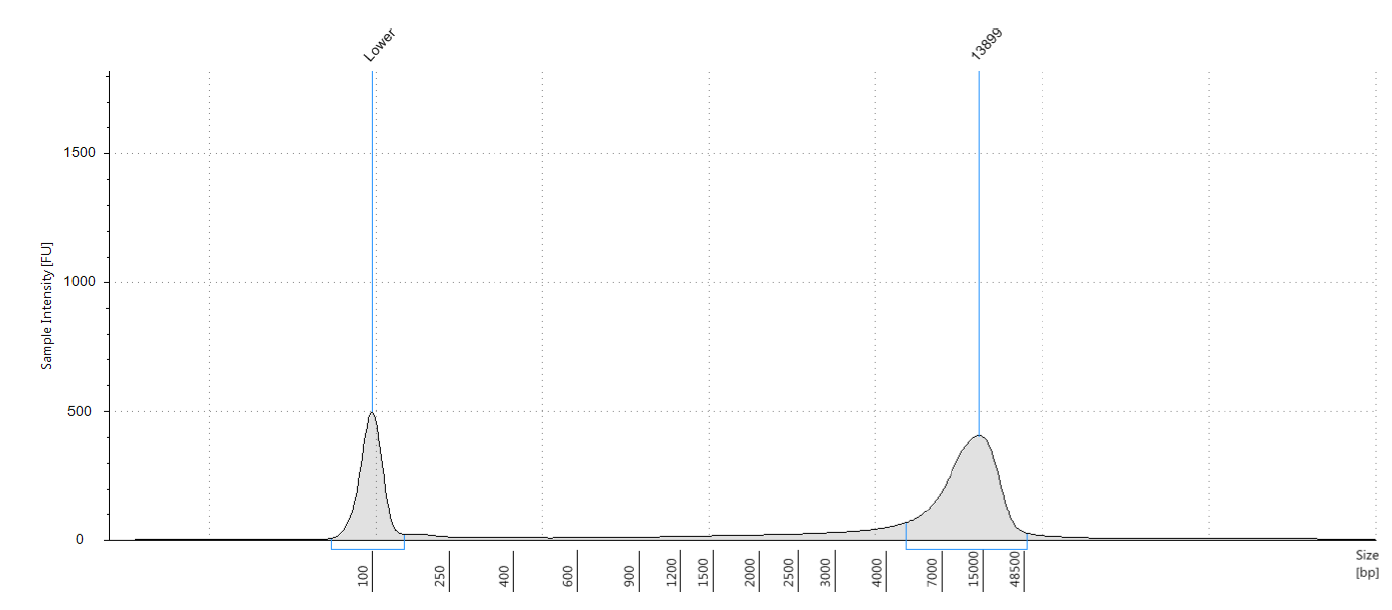

**Figure 2 Tapestation gDNA trace MoBio PowerSoil® for representative MiF (mixed forest) sample.**

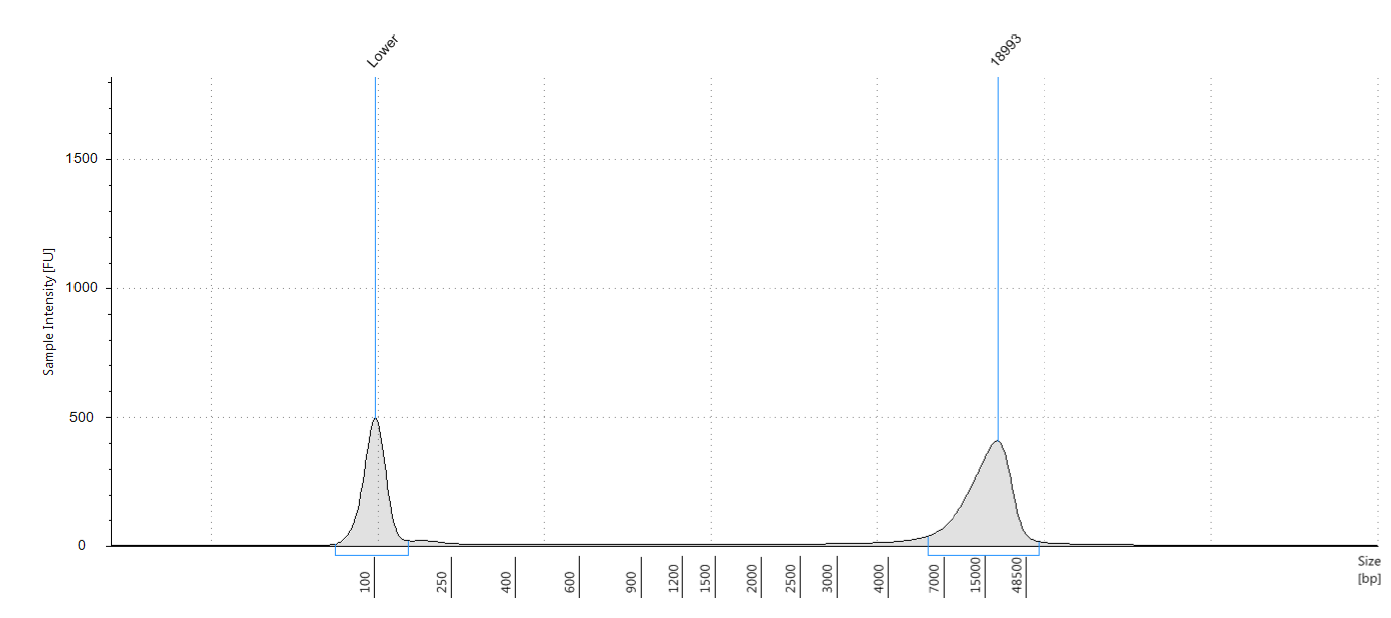

**Figure 3 Tapestation gDNA trace MoBio PowerSoil® for representative BrF (broad leafed forest) sample.**

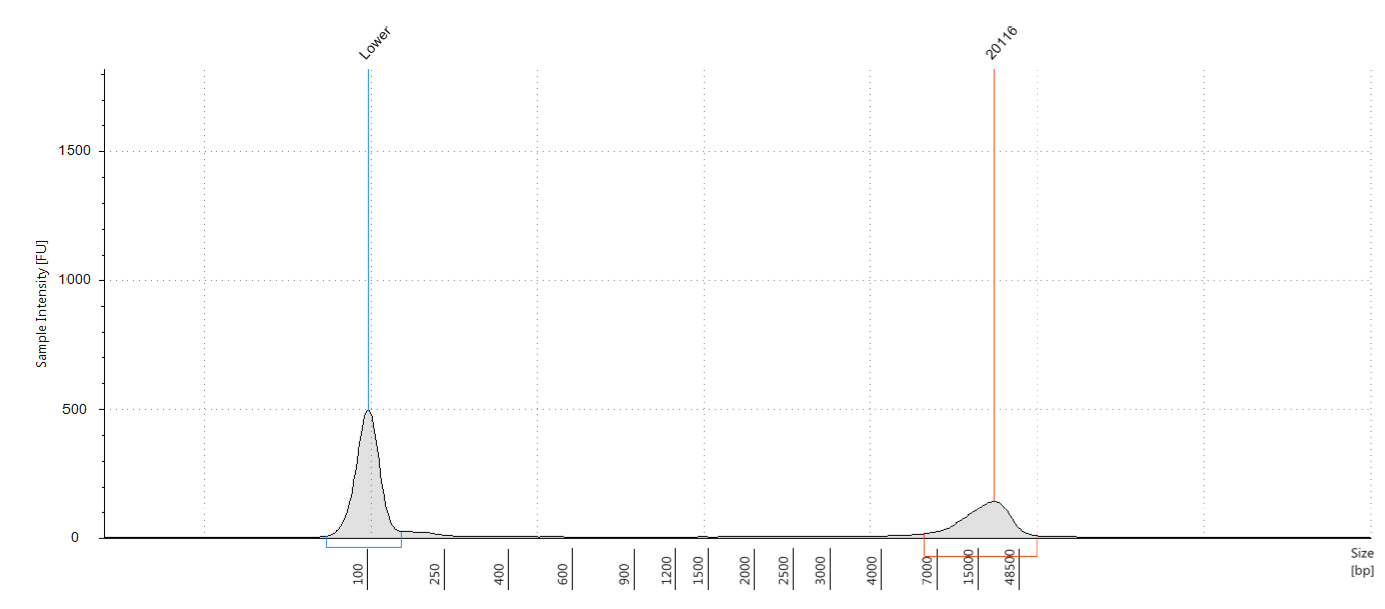

**Figure 4 Tapestation gDNA trace MoBio PowerSoil® for representative CoF (coniferous forest) sample.**

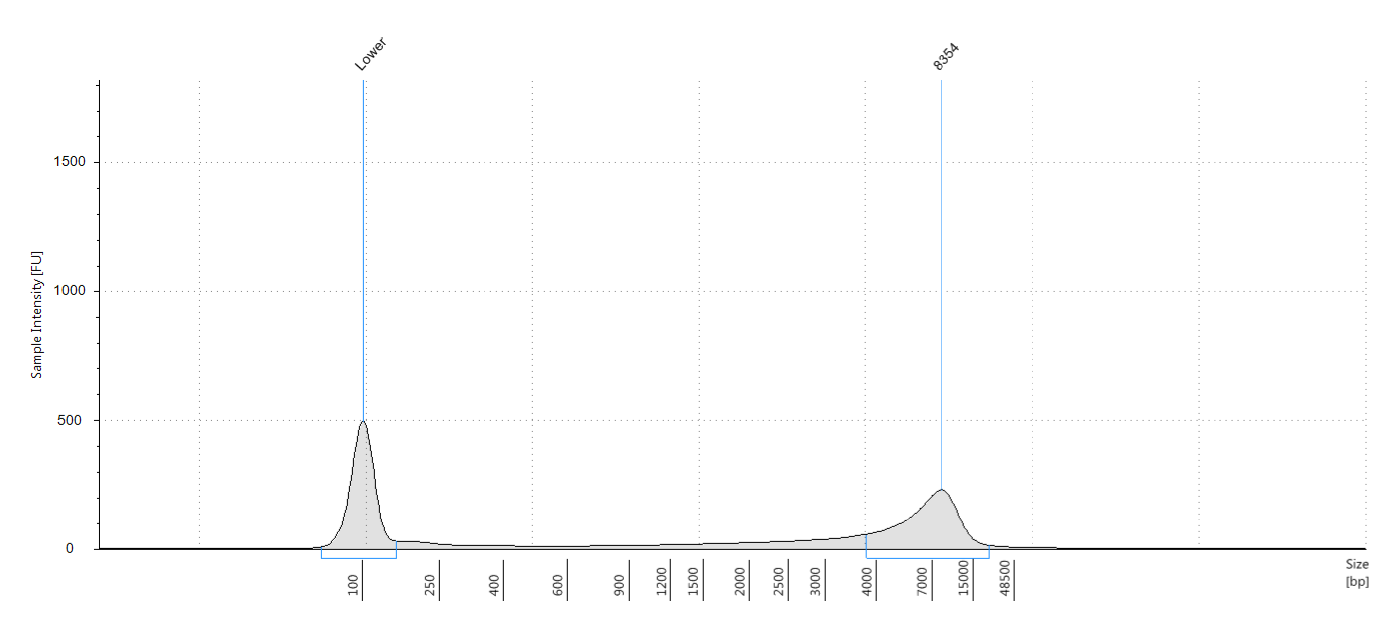

**Figure 5 Tapestation gDNA trace MP Biomedicals™ FastDNA™ SPIN kit for representative Cer (John Innes cereal compost mix) sample.**

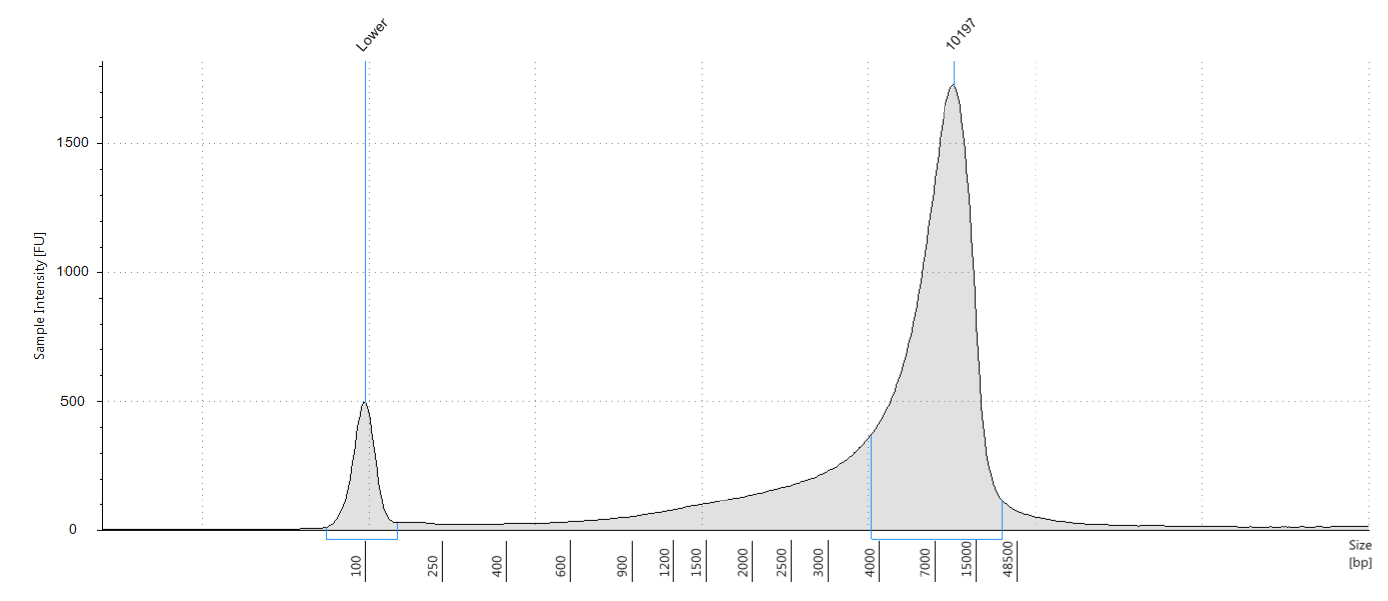

**Figure 6 Tapestation gDNA trace MP Biomedicals™ FastDNA™ SPIN kit for representative MiF (mixed forest) sample.**

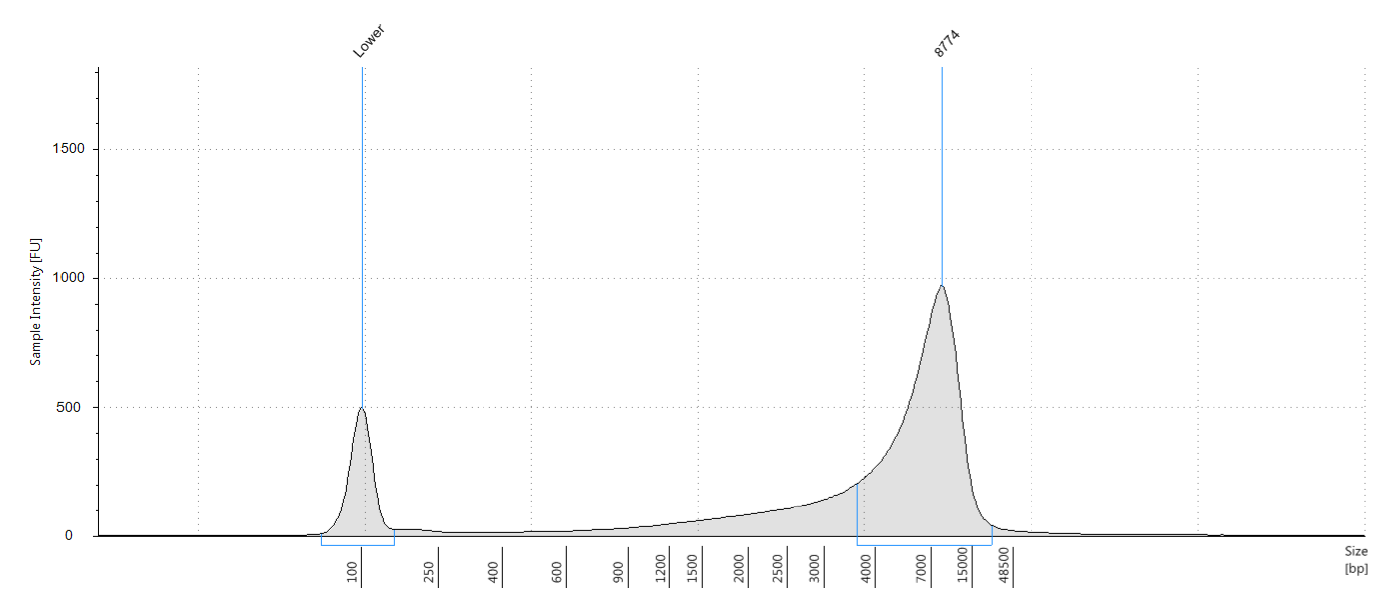

**Figure 7 Tapestation gDNA trace MP Biomedicals™ FastDNA™ SPIN kit for representative BrF (broad leafed forest) sample.**

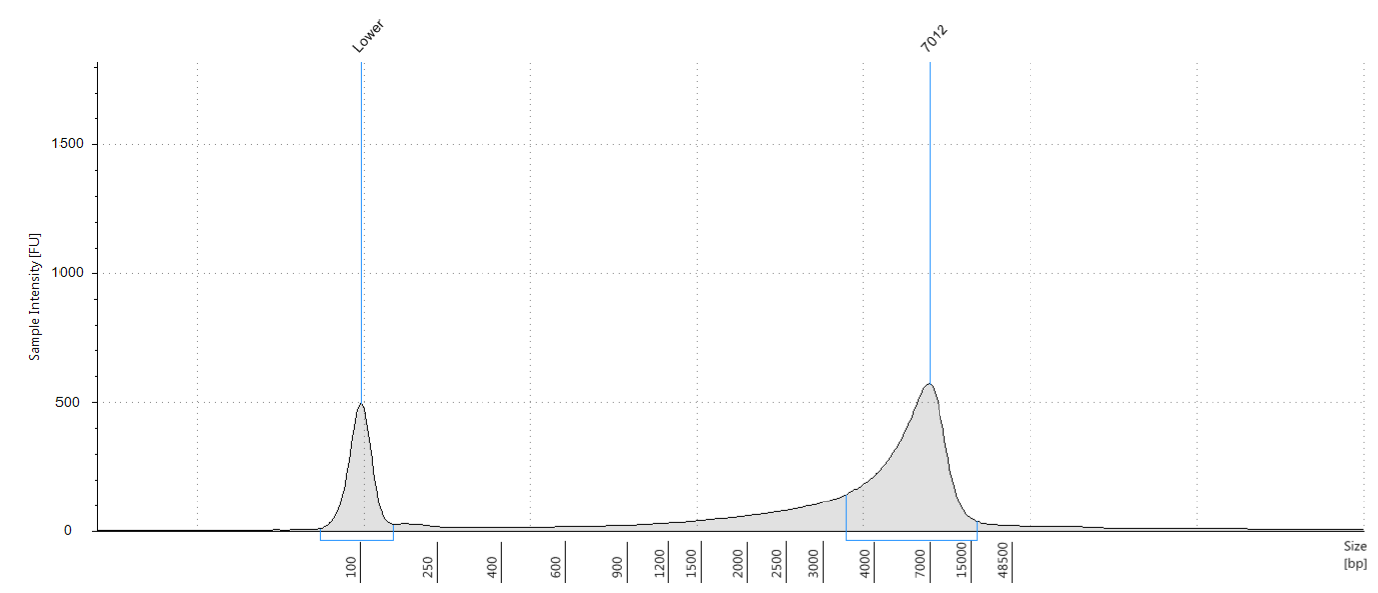

**Figure 8 Tapestation gDNA trace MP Biomedicals™ FastDNA™ SPIN kit for representative CoF (coniferour forest) sample.**

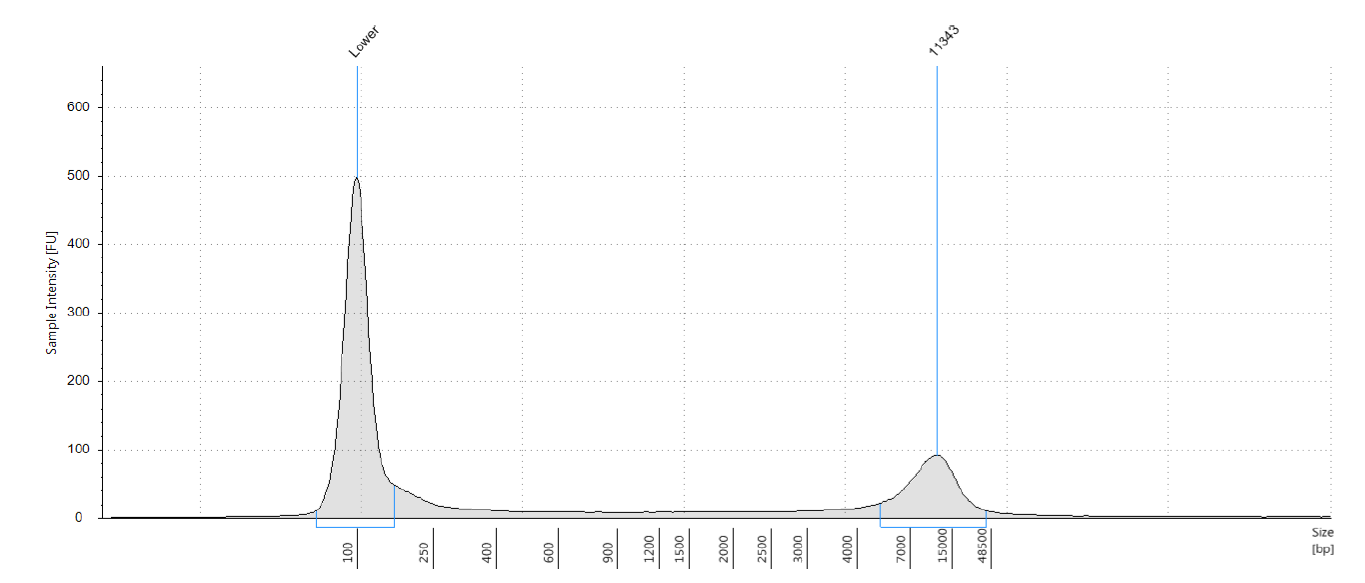

**Figure 9 Tapestation gDNA trace SDE method for representative CeR (John Innes cereal compost mix) sample.**

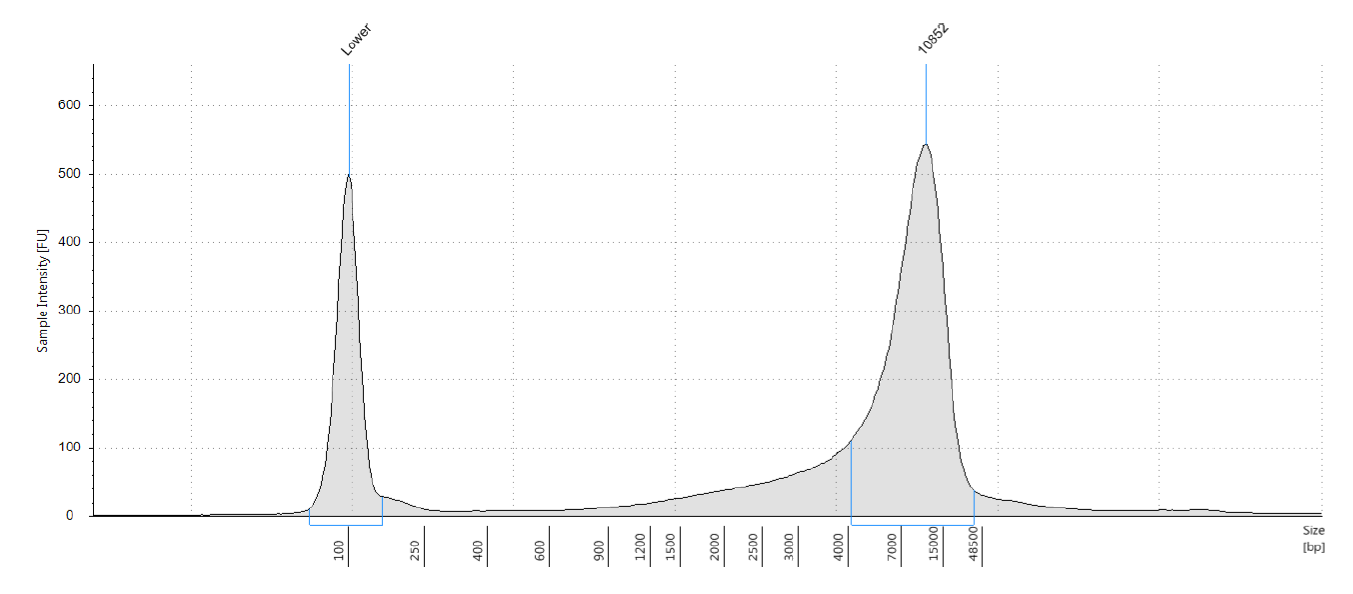

**Figure 10 Tapestation gDNA trace SDE method for representative MiF (mixed forest) sample.**

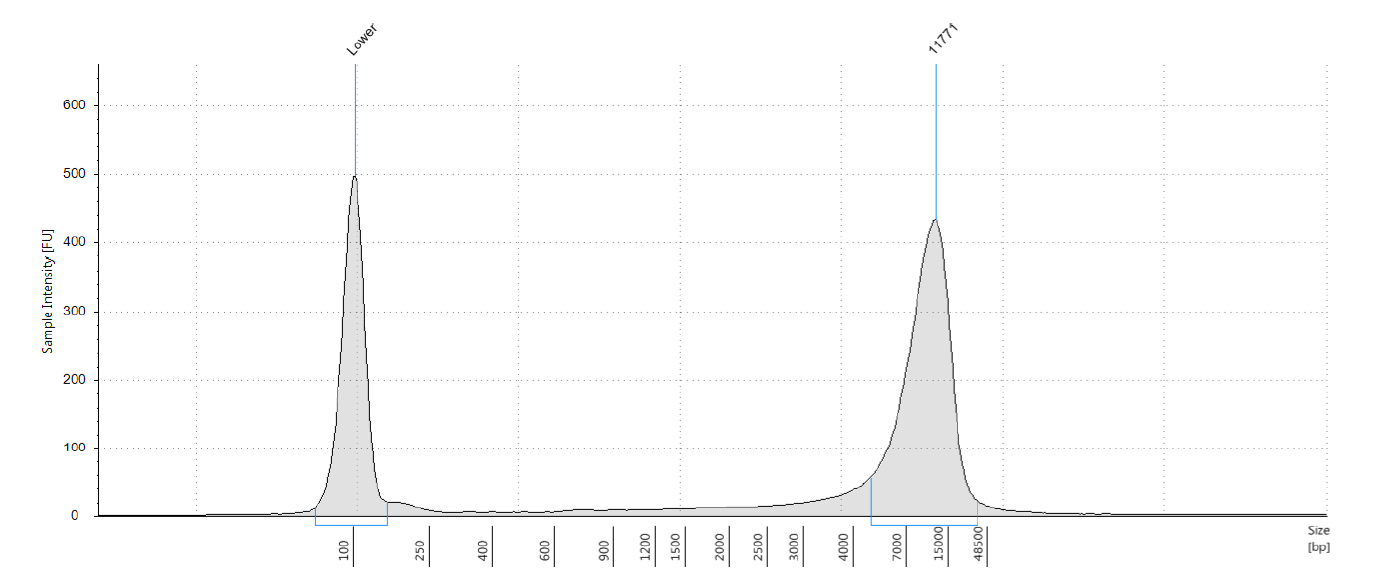

**Figure 11 Tapestation gDNA trace SDE method for representative BrF (broad leafed forest) sample.**

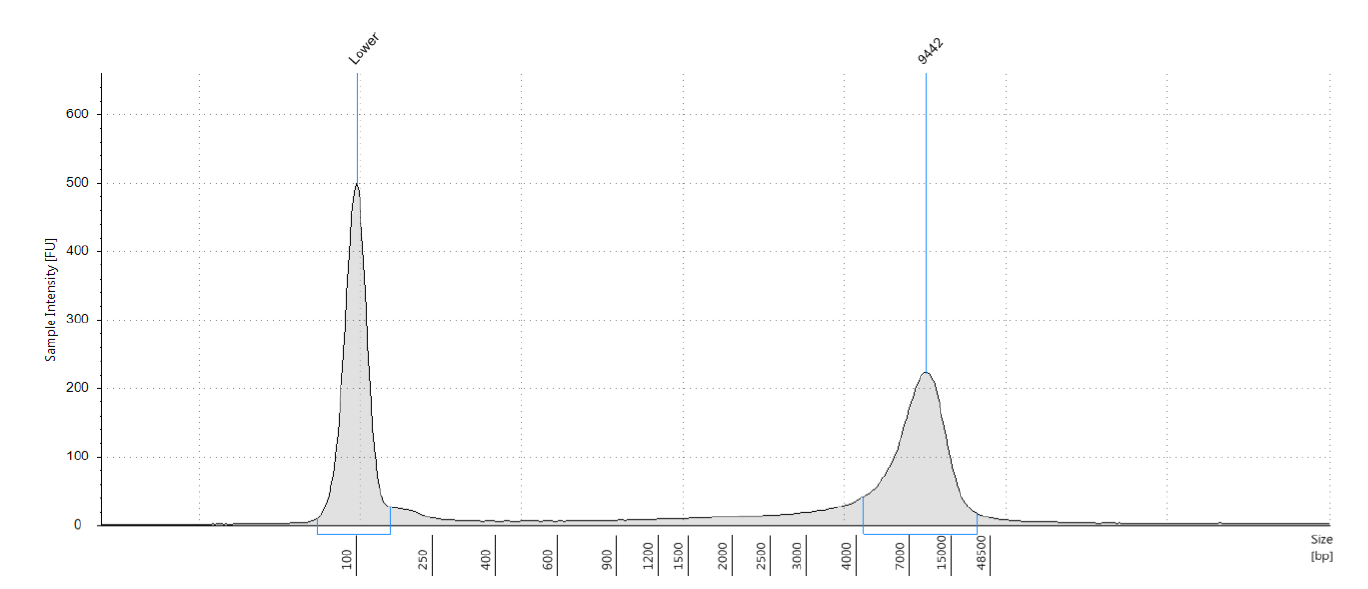

**Figure 12 Tapestation gDNA trace SDE method for representative CoF (coniferous forest) sample.**
