## Supplementary figures and images for "A low-cost pipeline for soil microbiome profiling"

### Additional_File_2

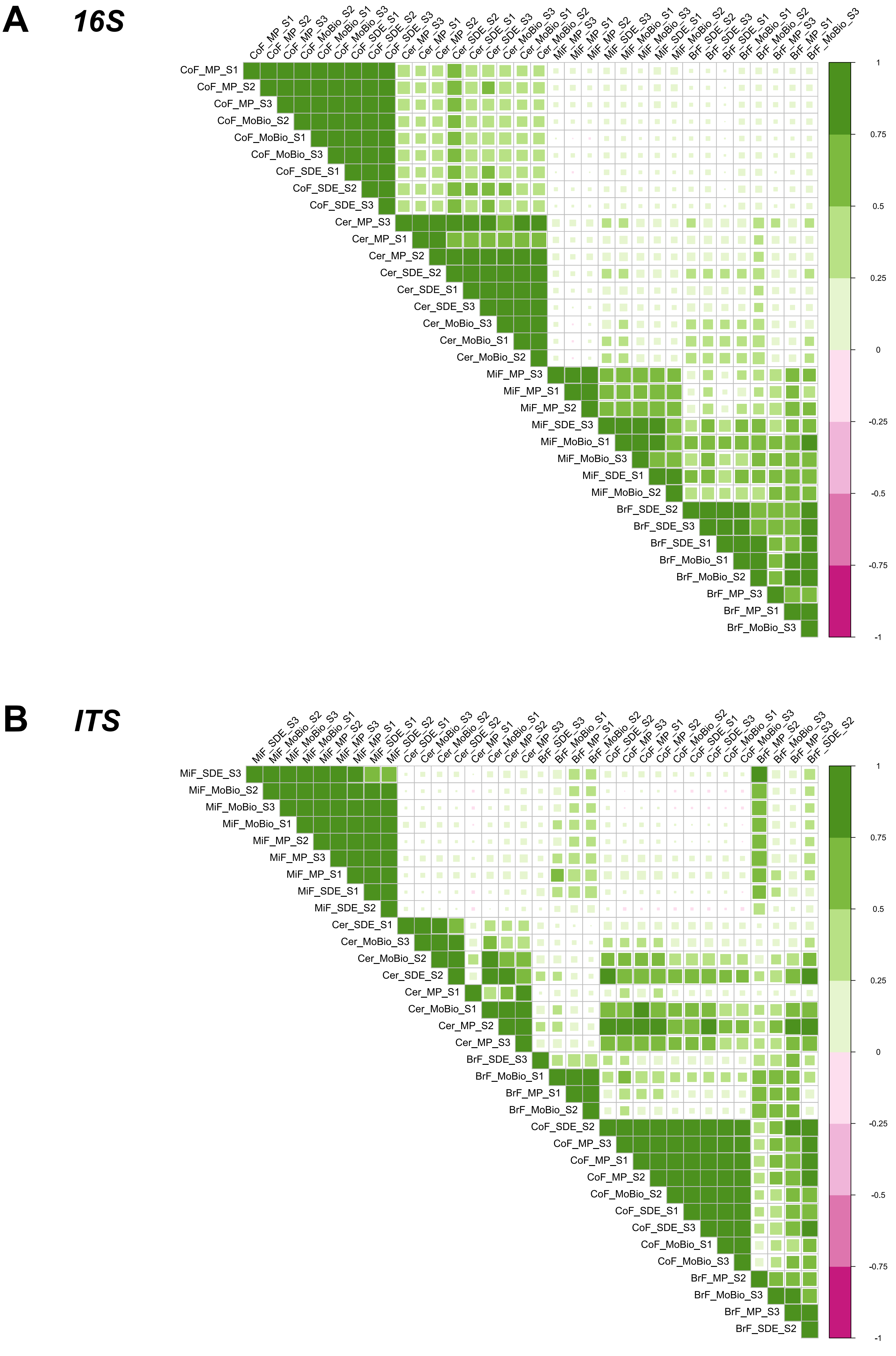

### Additional_File_3

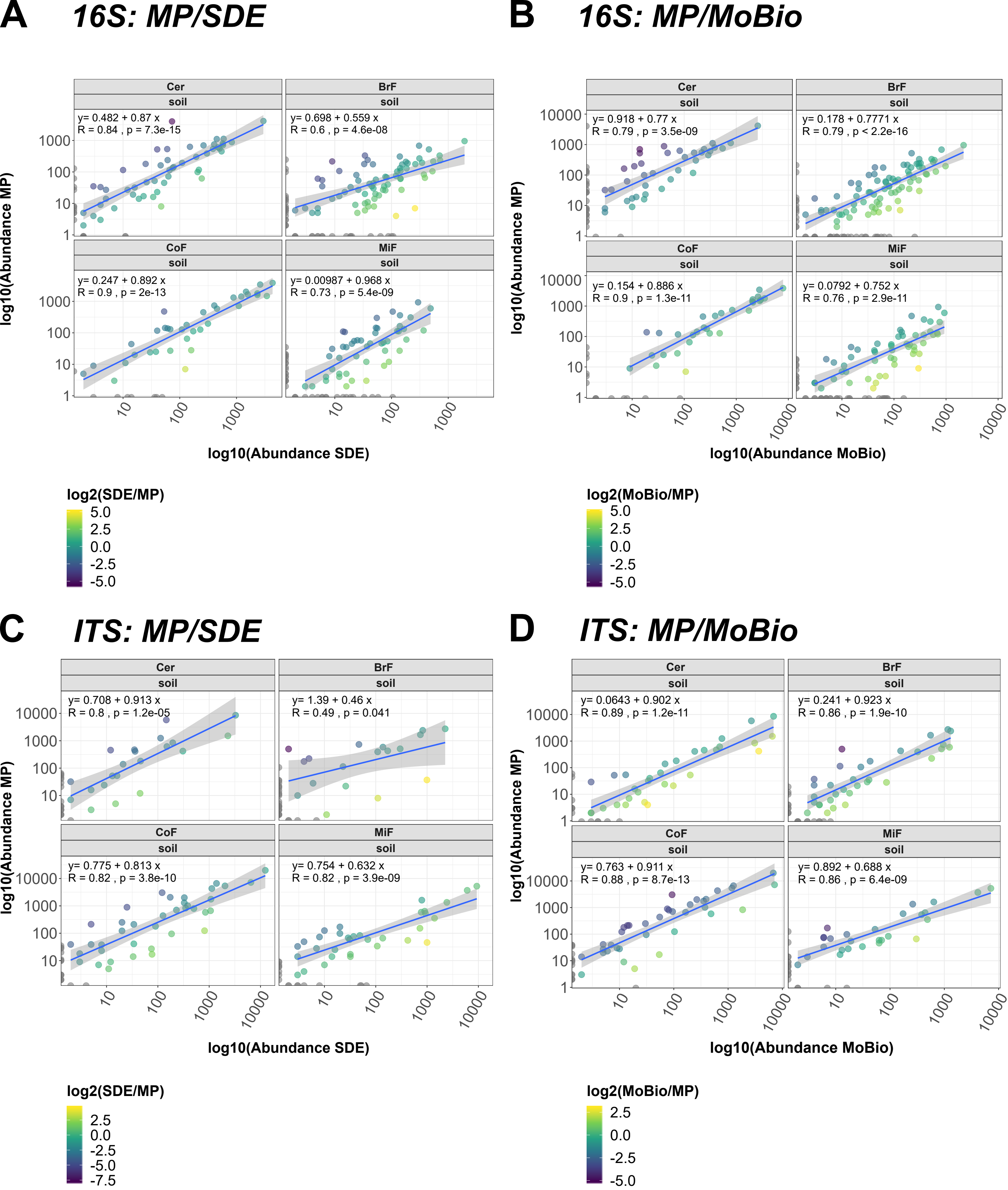
