## supplementalFigures for "A low-cost pipeline for soil microbiome profiling"

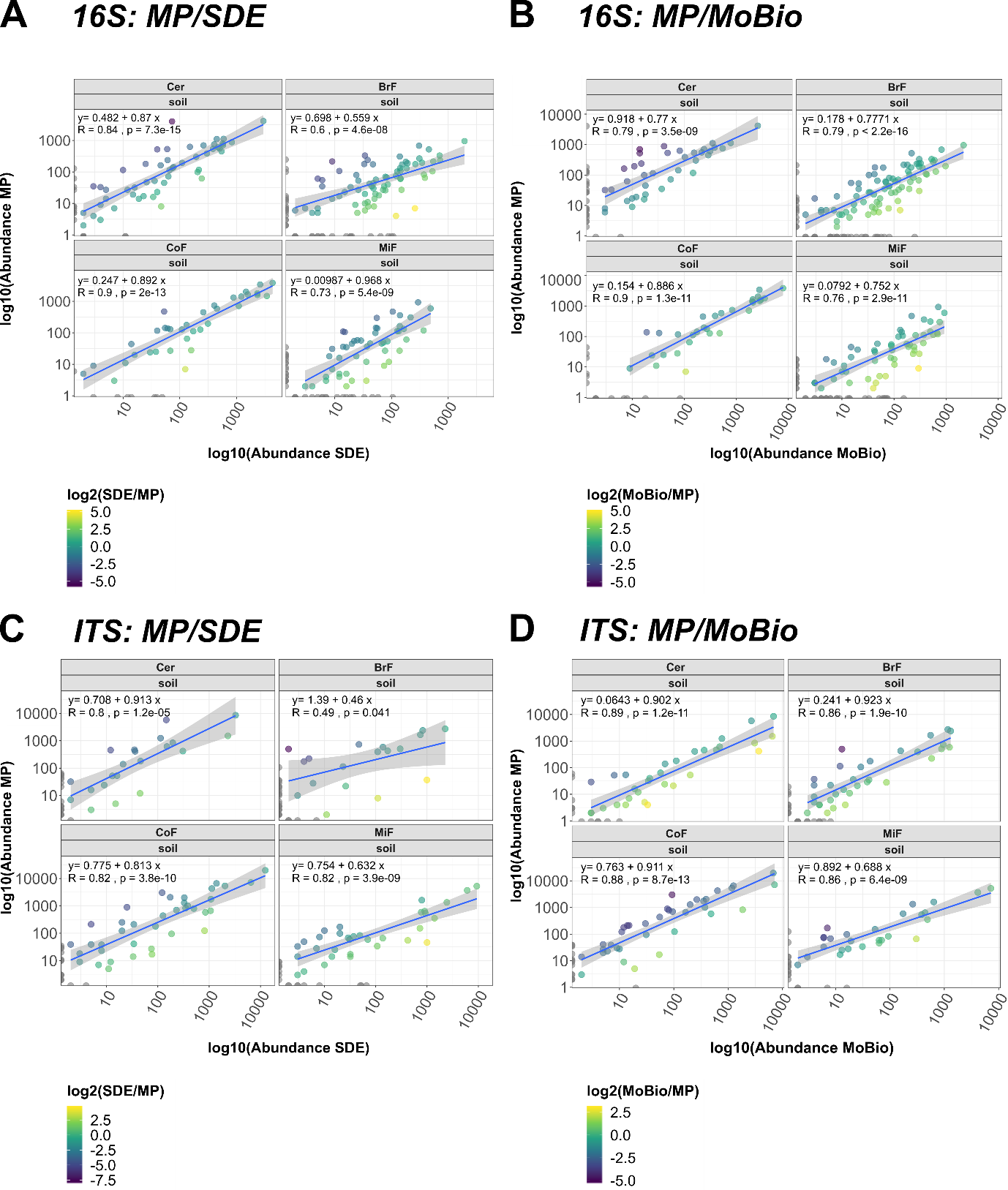

**Supplementary Figure 1** Correlation analysis SDE method versus MP Biomedicals™ FastDNA™ SPIN & MP Biomedicals™ FastDNA™ SPIN vs MoBio PowerSoil®(16S/ITS).

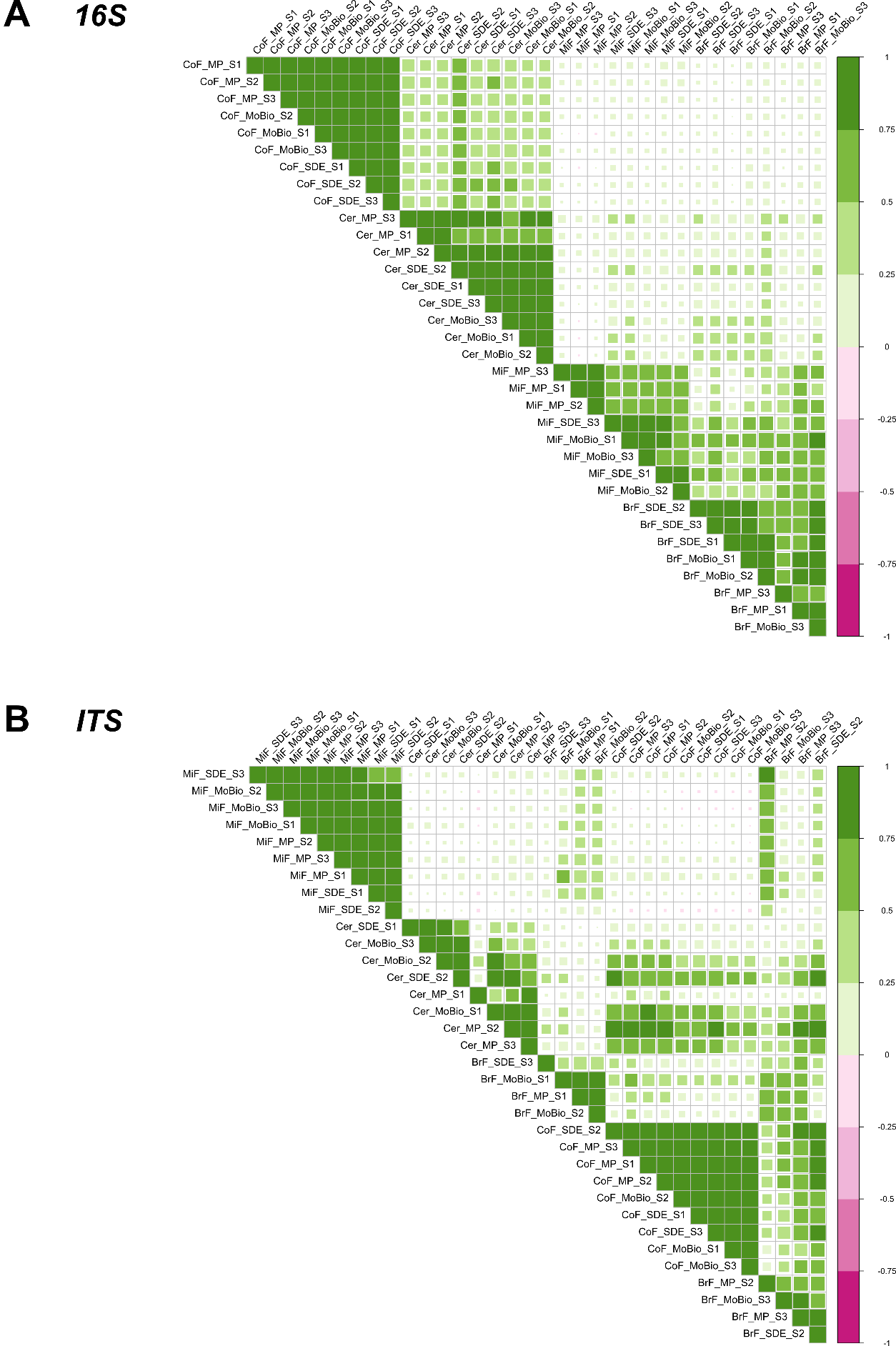

**Supplementary Figure 2** Corrplot of biological replicates of MP Biomedicals™ FastDNA™ SPIN, MoBio PowerSoil® and SDE methods. (**A**) shows bacterial correlation at Order level between MP Biomedicals™ FastDNA™ SPIN, MoBio PowerSoil® and SDE method considering biological replicates. (**B**) shows fungal correlation at Order level between MP Biomedicals™ FastDNA™ SPIN, MoBio PowerSoil® and SDE methods considering biological replicates.
